# Why motor learning involves multiple systems: an algorithmic perspective

**DOI:** 10.64898/2025.12.19.695526

**Authors:** Francesca Greenstreet, Jesse P. Geerts, Juan A. Gallego, Claudia Clopath

**Affiliations:** Department of Bioengineering, Imperial College London, London, UK; Neuroscience of Disease and Neuroscience Programs, and Centre for Restorative Neurotechnology, Champalimaud Foundation, Lisbon, Portugal

## Abstract

The initial stage of learning motor skills involves exploring vast action spaces, making it impractical to learn the value of every possible action independently. This poses a challenge for standard reinforcement learning approaches, which excel in constrained domains but struggle when the space of possible actions is high-dimensional. Recent work in machine learning has sought to mitigate this problem by combining deep reinforcement learning with a supervised learning system that reduces the complexity of the control policy space by learning low-dimensional embeddings of an action space. Here, we propose that in the mammalian brain, the cortico-cerebellar network learns these low-dimensional action embeddings in a supervised way, while the basal ganglia learn value and policies in this action embedding space using reinforcement learning. We trained this model on reaching tasks and show that, contrary to traditional models of the basal ganglia, it recapitulates features of neural activity whereby similar reaching movements are associated with similar neural activity patterns in the basal ganglia. We also demonstrate a link between learning these low-dimensional action embeddings and both generalisation and the limits of multi-task adaptation in human behavioural studies. Through this framework, we propose a novel computational view of how key motor regions of the brain interact to efficiently learn a new skill.

## Introduction

Everyday motor skills such as writing, driving and buttoning your jacket are not innate – they must be learned through trial and error in a process of skill acquisition and refinement. This process of motor skill learning involves using visual, and somatosensory feedback to learn to orchestrate the precise coordination of hundreds of joints and muscles while continuously evaluating success and adjusting for errors. Many regions of the brain have been shown to be involved in this learning process, most importantly the motor cortex, cerebellum and basal ganglia [1–4]. These three regions are heavily interconnected [2, 5–11], an anatomical observation that raises important functional questions: what drives the need for multiple regions in skill learning, and what is the nature of the resulting emergent function?

Individually, the motor cortex, cerebellum and basal ganglia have been hypothesised to each serve specific functions (Figure 1A). The basal ganglia receive dopaminergic reward prediction errors [12] and are thought to perform reinforcement learning [13–15]. The cerebellum and cortex both receive sensorimotor errors [16–18], which makes them ideal candidates for a supervised learning system [19, 20]. More specifically, the basal ganglia are thought to learn based on the predicted value of each action in a given state, which is then updated using reward information in the form of a dopaminergic reward prediction error, signalling the difference between the expected and obtained reward [12–15, 21, 22]. The cerebellum is key for motor adaptation [23, 24] as shown by the stark learning deficits observed in cerebellar patients [25–29], with the influential Marr-Albus-Ito model proposing that it operates as a supervised learning system where climbing fiber error signals train Purkinje cells to improve movement accuracy [30–33]. Even though the role of the motor cortex remains unclear [1], it has been extensively modelled as a supervised learning system [19, 34–38] that learns either directly based on sensorimotor errors or on cerebellar inputs reflecting sensorimotor errors [20, 39, 40].

**Figure 1:**
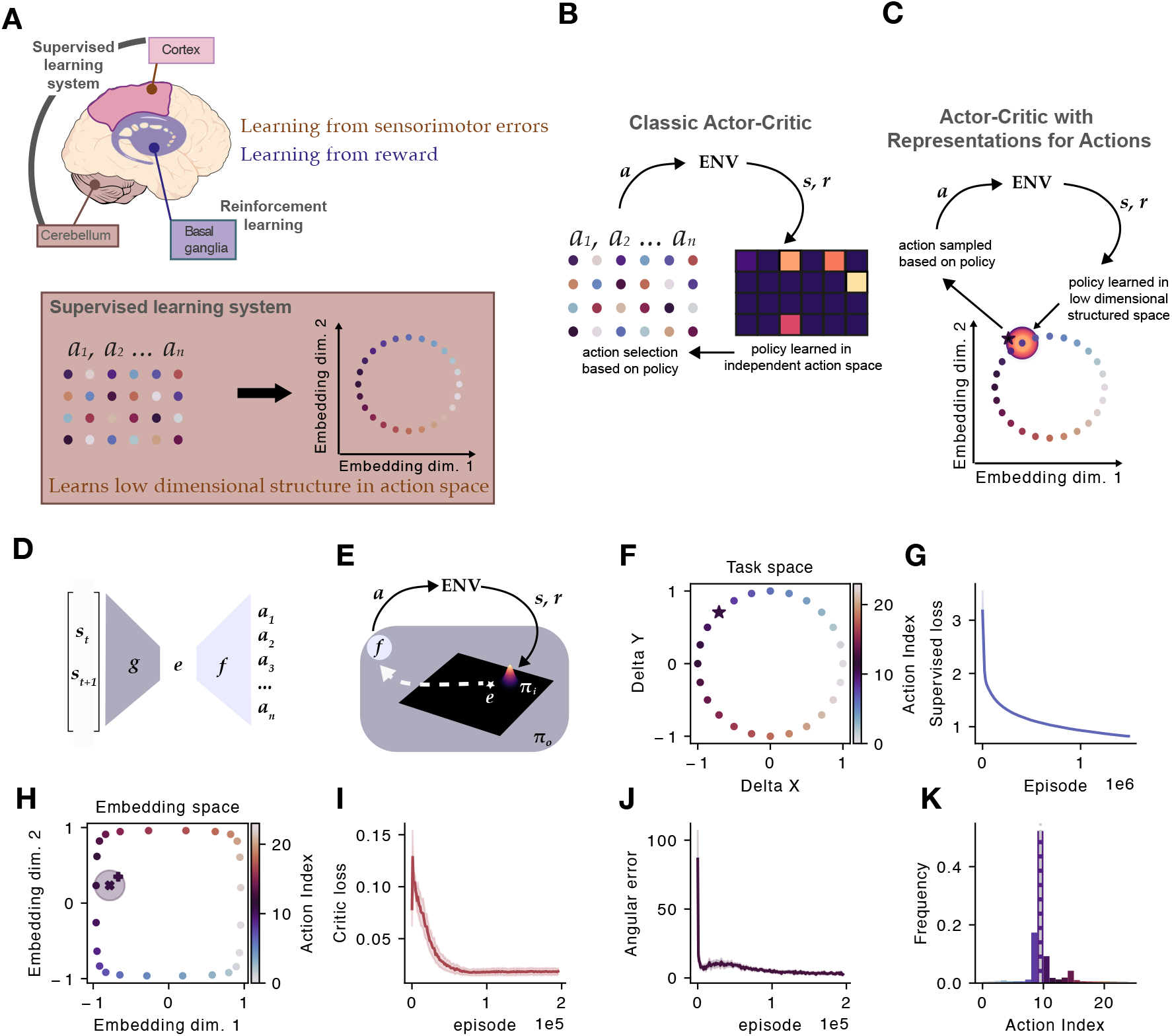
A multi-region model learns structured action representations. **(A)** The brain’s motor learning system involves multiple regions: the cortico-cerebellar network learns from sensorimotor errors (supervised learning), while the basal ganglia learn from reward (reinforcement learning). We propose that the supervised learning system learns a low-dimensional representation of actions, where similar actions are nearby in representation space. **(B)** In classic actor-critic reinforcement learning, actions are treated as independent. **(C)** In our model, the policy is learned in a low-dimensional structured embedding space, enabling generalisation. **(D)** Supervised model architecture: an encoder *g* maps state transitions (*s*_*t*_, *s*_*t*+1_) to embeddings *e*, which a decoder *f* maps to actions. **(E)** The agent learns an *internal policy π*_*i*_ over embeddings, with actions generated by decoding sampled embeddings via *f*. An outer policy *π*_*o*_ enables exploration. **(F)** Task illustration. centre-out reaching task with 24 movement directions arranged circularly in task space. **(G)** Supervised loss decreases during pre-training of the encoder-decoder network. **(H)** Learned embedding space preserves the circular structure of the task. Each point indicates the model’s middle layer activity in response to a particular (*s*_*t*_, *s*_*t*+1_) pair. The *×* with shaded area represents the agent’s learned internal policy, and the + indicates a single sample from the policy. **(I)** Critic loss (squared TD error) and **(J)** angular error decrease during reinforcement learning, demonstrating successful policy learning. **(K)** Learned action distribution. Dashed line shows optimal action. Shaded error bars represent SEM across 10 random seeds in **G, I, J**.

Despite substantial anatomical evidence that the basal ganglia, cortex and cerebellum are highly interconnected [2, 5–10], there are few algorithmic-level theories for what purpose these connections serve, that is, how the cortico-cerebellar supervised learning should relate to reinforcement learning in the basal ganglia. Our goal here is to develop a computational-level theory for the nature of these interactions, and how sensorimotor error driven supervised learning in cerebellum and cortex relate to reinforcement learning in the basal ganglia.

In machine learning, there is a growing body of literature showing that reinforcement learning, which is typically slow [41, 42], can benefit from supervised learning to reduce the dimensionality of the action space that value needs to be learned over [43–48]. This is particularly relevant for real-world reinforcement learning problems in robotics, where the dimensionality of the action space is typically large (i.e. all combinations of joint movements for a robotic arm), rendering classical reinforcement learning approaches inefficient [46, 48]. These models combine supervised and reinforcement learning to leverage the insight that actions are not independent, but rather they are organised in a structured, typically low dimensional, way (Figure 1B) –for example, in robotics there are multiple joint configurations that can achieve the same end-effector position or movement outcome. That way, these so-called Reinforcement Learning with Representations for Actions [43] models use supervised learning to train a dedicated neural network to predict the action that caused the transition from one state to the next^1^. This supervised network reduces the dimensionality of the action space from the number of possible actions to a lower dimensional latent embedding space. A separate network can be trained to learn a policy directly in this “action embedding space” (Figure 1C). This approach drastically speeds up learning by improving how the space of potential actions is explored, and leads to better alignment between action space and the task [43, 46].

Here, we take inspiration from these machine learning models and propose that the brain may operate similarly, with supervised learning systems (motor cortex and cerebellum) providing structured representations of the action space that facilitate more efficient reinforcement learning in the basal ganglia. This framework offers a principled explanation for why multiple, interacting learning systems might be advantageous: rather than learning values over the full high-dimensional action space, the basal ganglia could learn more efficiently in a lower-dimensional space structured by the supervised learning system. We test this proposal by training an action-embedding model on a standard centre-out reaching task [52, 53] and demonstrate that learning policies in the embedding action space naturally leads to structure between policies that is consistent with the structure seen in recent neural recordings from the basal ganglia [54]. Furthermore, we show that this framework enables flexible generalisation in the action space and can account for key phenomena in motor adaptation, including generalisation to similar actions [55] and experimentally observed limits of adapting to multiple visuomotor rotations simultaneously [56]. Taken together, our results suggest that the interacting brain regions involved in motor learning may reflect a modular architecture that leverages learned structure in the action space for more efficient learning.

## Results

### Model overview

Here, we introduce a computational model of cerebellar-cortico-basal ganglia interactions that integrates a corticocerebellar module, performing supervised learning [19, 34–38], and a basal ganglia module performing reinforcement learning [13–15]. Our hypothesis is that organising reinforcement learning policies in a way that policies for actions that share similar consequences are close together will lead to two key benefits for motor learning: faster learning and generalisation between similar actions.

We tested this hypothesis by building a model in which the corticocerebellar module learns a low-dimensional embedding of a high-dimensional action space in a supervised manner. This learned low-dimensional embedding captures a similarity structure between actions that share similar consequences. The basal ganglia reinforcement learning module, in turn, outputs its policy directly in this learned embedding space.

More precisely, the supervised learning module, proposed to be implemented by the cerebral cortex and cerebellum, takes the form of an encoder-decoder network, trained with a cross-entropy loss (a standard measure of how well the network’s predictions match the actual actions) and backpropagation (Figure 1D). The action encoder, *g*, is a neural network that first maps state transitions (*S*_*t*_ and *S*_*t*+1_) to a low-dimensional embedding space, *e*. The action decoder, *f*, then linearly maps this embedding from the latent action space to the output action space, 𝒜. We refer to the relationship between a point in embedding space and the output action space as the action-embedding mapping. The role of this encoder-decoder network is to predict the action that was taken given the current and next state, *P* (*A*_*t*_|*S*_*t*_, *S*_*t*+1_). The supervised loss is calculated given the predicted (*Â*_*t*_) and actual action (*A*_*t*_) and serves to update the network weights to improve action prediction in subsequent trials. This way, the model learns to output low-dimensional embeddings that capture the similarity structure of actions: actions with similar movement consequences map onto nearby parts of the action embedding space (Figure 1B).

In contrast to classic reinforcement learning models where policies are learned directly over the actions, our reinforcement learning module learns “internal policies” directly over the action embedding space, *e* (Figure 1C). Points in the embedding space are then sampled from the policy and subsequently mapped to the action space using the action mapping learned by the supervised network (Figure 1E). Thus, policies are learned in the embedding space, but points sampled from this space can be translated back to the action space in order for the agent (or animal) to act. Crucially, the “translator” is the action decoder (*f*) from the supervised network, which ensures that the policies are compatible with the structure of the embedding space learned by the encoder-decoder network.

The reinforcement learning module that serves as an idealisation of the basal ganglia network takes the form of an actor-critic architecture [57] which, unlike the supervised cortico-cerebellar network, is trained with a temporal-difference reward prediction error learning signal. Thus, the model harnesses fast supervised learning to learn a structured representation of possible actions, which in turn speeds up reinforcement learning, particularly as the size of the action space increases [43] (Supplemental Figure A.1).

### Similar actions have closer policies in embedding space

We first tested whether representations for similar actions are organised close together in the embedding space by comparing the activity of the model to that of the mouse basal ganglia during the same reaching task in mice [54] (Figure 1F). Agents were trained to reach from the centre to a target zone in order to get a reward. First, we trained the supervised module to learn representations with the agents following a random policy. Over training, the network became better at predicting which action caused each observed state transition, as indicated by the decreasing classification loss (Figure 1G). That is, it learned meaningful relationships between actions and their consequences in the environment.

We visualised the structure of the embedding space by passing the state transitions that would be caused by taking each available action through the first part of the supervised network, *g*. As predicted, the structure of the action space (Figure 1F) was preserved in the embedding space (Figure 1H), with actions that lead to neighbouring endpoints having neighbouring embeddings (Supplementary Figures A.2 and A.3). We then trained the actor-critic network to learn an internal policy in the embedding space with the aim to maximise reward (Figure 1H shows an example learned Gaussian policy). The agent learned to predict rewards, shown by the critic’s loss approaching zero (Figure 1I) and the selected action tending towards the optimal one (Figure 1J & K). This confirmed that our supervised network learned functionally relevant structure in the consequences of all available actions, and that the reinforcement learning network could exploit this structure to successfully take the appropriate action given the state of the agent and its goal.

We next compared our model to experimental recordings of neural activity. Classical models of motor control propose a division of labour: cortex specifies continuous movement parameters while basal ganglia select among discrete action options (Figure 2A). Under this view, activity in the basal ganglia should be distinct for different actions regardless of their similarity – a prediction captured by selection encoding models (Figure 2D, left), and consistent with previous studies showing very different basal ganglia activity in animals trained to respond with two distinct actions, such as left/right turning [58–60], a left/right/up/down choice [61], or a push/pull task [62]. Our model makes a different prediction: because the basal ganglia learn policies in a structured embedding space where similar actions are nearby (Figure 2B), striatal activity patterns should reflect action similarity, with similar actions being associated with overlapping activity patterns. Recent work supports this prediction: Park et al. [54] showed that striatal dynamics underpinning reaches to close-together targets are highly similar – to an extent comparable to motor cortex – and reflect time-varying movement kinematics (Figure 2D, middle).

**Figure 2:**
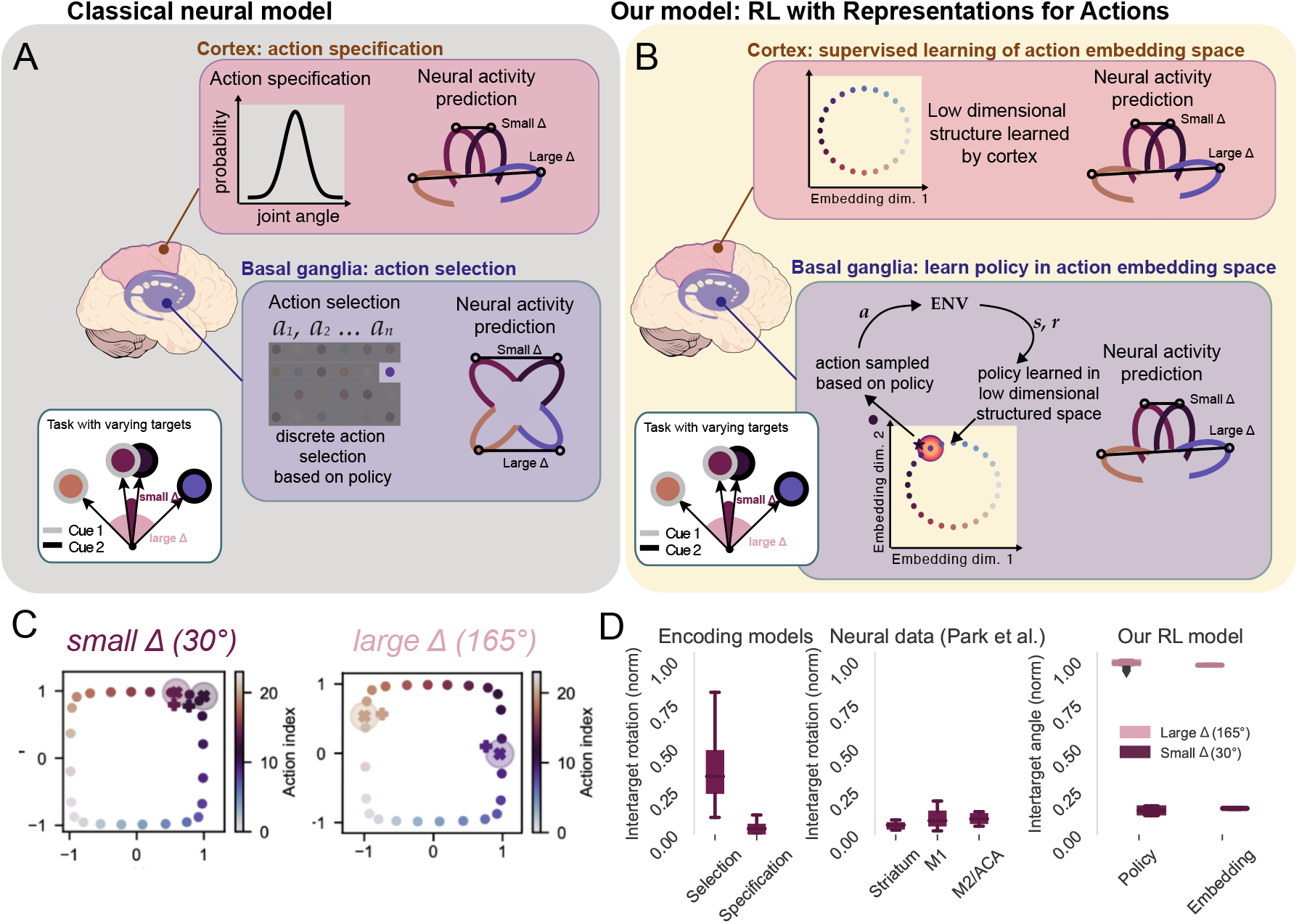
Our model predicts specification-like neural activity in the basal ganglia, consistent with experimental data. **(A)** Classical models propose that cortex performs action specification (continuous parameterisation) while basal ganglia perform action selection (discrete choices), predicting distinct basal ganglia representations regardless of action similarity. **(B)** In our model, cortex learns a structured action embedding space where similar actions are nearby, and basal ganglia learn policies in this space, predicting that basal ganglia representations should reflect action similarity. **(C)** Learned policies for small (30°, left) and large (165°, right) target separations. Policies overlap for similar targets and diverge for dissimilar targets. **(D)** Intertarget representational distance. Selection encoding models [replotted from 63] predict high distances regardless of target similarity; specification models predict low distances for similar targets (left). Neural data from striatum, M1, and M2 show specification-like patterns (middle; data replotted from [54]). Our model reproduces this: low distances for small Δ, high for large Δ (right).

To test whether our model recapitulates these experimental observations, we simulated reaches to different pairs of targets, placed either close together (small Δ = 30°) as in Park et al. [54], or far apart (large Δ = 165°) (Figure 2C). The model learned both tasks successfully (Supplementary Figures A.4 and A.5). As predicted, policies for nearby targets overlapped substantially in embedding space, while policies for distant targets occupied distinct regions (Figure 2C). We quantified this by computing the normalised angle between policy means across ten random seeds, finding low angles for small Δ and high angles for large Δ (Figure 2D, right)—closely matching the specification-like pattern observed in striatum (Figure 2D, middle). Critically, unlike the encoding models used by Park et al. which assume a fixed representational structure, our model learns this structure through interaction with the environment. This demonstrates that action specification-like activity patterns in the basal ganglia can emerge naturally from reinforcement learning in a structured action space where policies are learned in a low-dimensional embedding space that preserves action similarity, allowing efficient generalisation between related movements.

### Supervised embedding of similar actions enables flexible generalisation

The results above are consistent with the view that the brain learns a structured embedding of actions that it uses to perform reinforcement learning. This raises the question of what functional purpose might learning policies over such an embedding serve, as opposed to learning directly in the action space? First, learning with such a representation speeds up learning when there are many possible actions [43]. But this is not the only advantage of learning structured action representations: meaningful representations of *states* can aid generalisation between similar states [64]. Therefore, reinforcement learning using structured action representation may enable generalisation between similar *actions*. As actions that cause more similar state transitions are close to each other in the action embedding space, learning a policy that samples from this embedding space should lead to generalisation over actions with nearby embeddings.

We investigated this generalisation phenomenon in the context of visuomotor rotation experiments. In this paradigm, participants performing in a visually-guided reaching task, are subjected to a rotation of the visual feedback about the position of the cursor by a fixed angle around the centre of the workspace (Figure 3A) [1, 55]. Humans rapidly correct for this visuomotor perturbation within a few tens of trials in a process known to involve the cerebellum [e.g. 26–29, 65]. Interestingly, if humans are tested on targets other than the one used for the initial adaptation (Figure 3A), they show a characteristic pattern of generalisation of the learned correction (Figure 3B, adapted from [55]): generalisation is greatest around the trained target and drops off as test targets are placed at angles further from the trained target [66–69].

**Figure 3:**
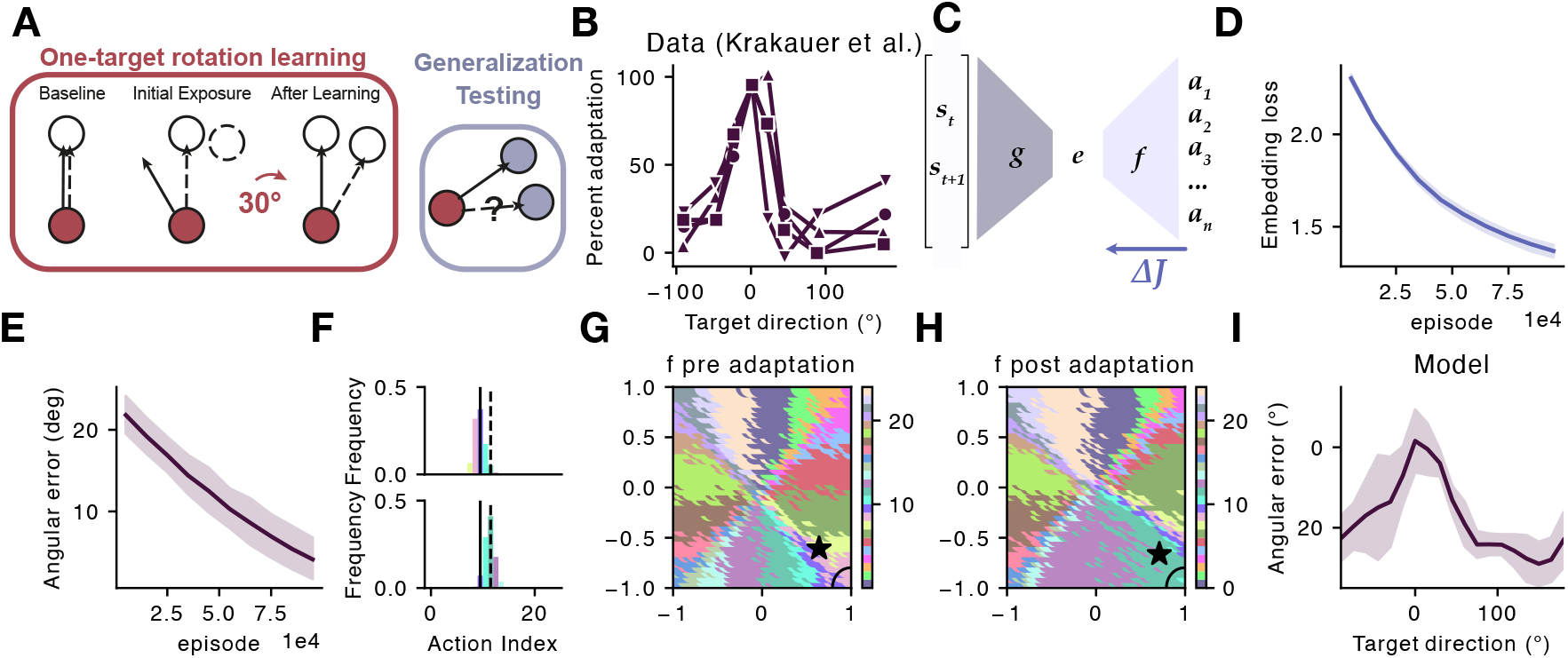
Learning action embeddings enables human-like generalisation in visuomotor adaptation. **(A)** Visuomotor rotation paradigm: during baseline, visual feedback matches movement direction; during initial exposure to a 30° rotation, reaches miss the target; after learning, reaches compensate for the rotation. Generalisation is tested by probing adaptation at untrained target directions. **(B)** Human data show characteristic local generalisation: adaptation transfers strongly to nearby untrained targets but decreases with angular distance from the trained target (data replotted from Krakauer et al., 2000). **(C)** We model adaptation by retraining only the decoder *f* while freezing the encoder *g* and RL components, representing fast supervised learning from sensorimotor errors. **(D)** Embedding loss decreases during adaptation training. **(E)** Angular error decreases as the model learns to compensate for the rotation. **(F)** Action selection before (top) and after (bottom) adaptation, showing a shift toward the compensatory action. **(G–H)** Decoder mapping from embedding space to actions before **(G)** and after **(H)** adaptation. The mapping rotates locally around the region where the policy is concentrated (star), while remaining largely unchanged elsewhere. **(I)** The model exhibits human-like generalisation: strong transfer to nearby angles with distance-dependent falloff, emerging naturally because decoder plasticity is greatest where the policy samples most frequently.

To model this behaviour, we used a network that had been pre-trained on a single target reaching task (as in Figure 1). This pre-training captures the baseline reaches, prior to the visuomotor rotation being introduced. For the reinforcement learning system, this means learning a policy through a process of exploration and reward-based feedback. Following this initial training, we replicated the visuomotor rotation paradigm by keeping the same target while rotating visual feedback by 30°, which the network experienced as the original actions leading to a different observation shifted by 30°. As motor adaptation is a fast cerebellar-dependent process based on sensorimotor errors [29] that is not disrupted by basal ganglia dysfunction according to studies in Parkinsonian patients [70–72], we modelled this fast adaptation process as happening through supervised learning in the corticocerebellar network, rather than through reinforcement learning in the basal ganglia, which is slow [41]. Moreover, since the adaptation experiment consists simply of a rotation, our model does not need to re-learn its action embedding structure. Instead, it should suffice to simply re-learn the action decoder weights, *f*, which linearly map the selected policy onto its associated action.

We tested this prediction in our model by freezing the weights in the reinforcement learning network and the action encoder network, *g*; therefore, only the weights in the action decoder network, *f*, were retrained during the adaptation phase (Figure 3C). As expected, during the first trials after the rotation was included, the same action led to an unexpected state transition, causing a large discrepancy between the action predicted by the supervised network and the action taken. As *f* learned to predict the new actions mapped to the state transitions, both the supervised error of the corticocerebellar network (Figure 3D) and the angular reach error of the agent decreased (Figure 3E). This led to a shift in the action distribution (Figure 3F) to be centered around the new optimal action, recapitulating the behaviour of humans performing the same one-target adaptation visuomotor adaptation task [55]. This learning was mirrored in the action embedding space, where a rotation was seen locally around the policy mean (Figure 3H) when compared to the original embedding-action mapping (Figure 3G).

This local rotation of the embedding-action mapping suggested that the agents might show similar generalisation properties to humans (Figure 3B). We tested this by freezing all the weights and evaluating policies for each relevant location in the embedding space. As seen in humans [55, 66–69], our models generalised rotation adaptation to nearby targets but not to distant targets (Figure 3I), suggesting that fast relearning of an embedding-action mapping can account for human adaptation generalisation behaviour. This generalisation profile was consistent across all 24 initial learning angles (Supplementary Figure A.6).

### A shared embedding-action mapping explains dual adaptation limits

As well as being able to generalise adaptation across similar actions, humans also can simultaneously adapt to multiple visuomotor rotations if the targets are sufficiently distant from one another [56]. In our model, visuomotor adaptation is driven by locally relearning the embedding-action mapping for embeddings close to the original policy mean. Thus, the embedding-action mappings are most altered for actions similar to the adapted action (Figure 3H & I). This led us to predict that performance at simultaneous learning of multiple visuomotor rotations would depend on the proximity of the reach targets, consistent with human experiments [56]. This prediction is a direct consequence of our model as local relearning of the embedding-action mapping for two reach angles would interfere if the targets are close together, but not if they are far apart.

To test this, we simulated the experiment by Woolley et al. [56] in which humans were given visuomotor rotations of 30° clockwise or 30° counterclockwise to two targets that either lay on top of each other in the visual space (indistinguishable) or targets that were directly opposite each other on the reach circle (Figure 4A-B). Humans are able to learn both clockwise and counterclockwise rotations in the case of the two opposite targets but not if the targets are the same.

**Figure 4:**
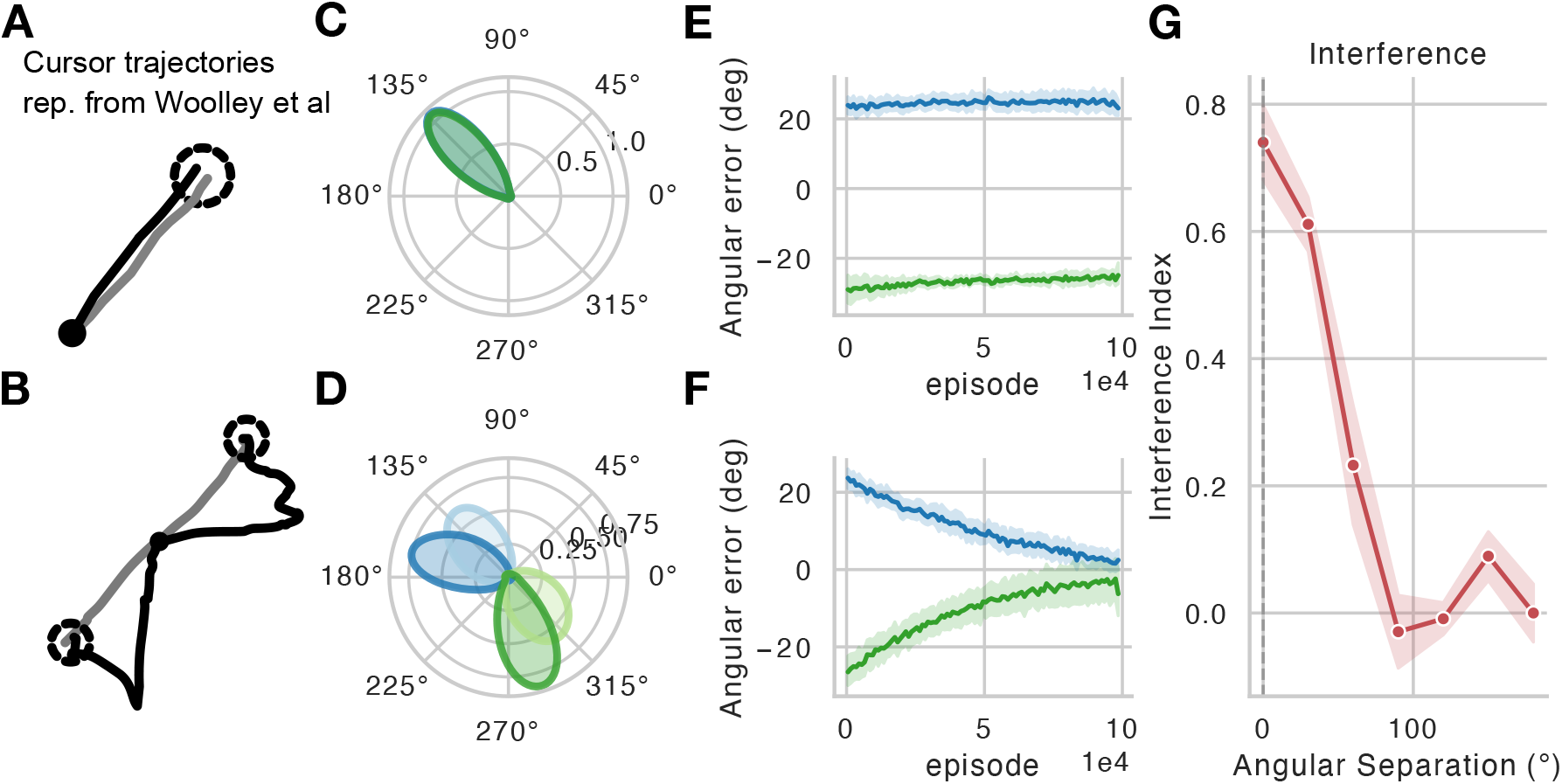
Dual adaptation to opposing visuomotor rotations. **(A–B)** Humans can learn opposing visuomotor rotations (+30 and −30) when applied to targets far apart **(B)**, but show interference when rotations are applied to nearby targets **(A).** Trajectories re-drawn from [56]. **(C– D)** Nearby targets have overlapping policies in embedding space **(C)**, forcing competing changes to the same decoder region. Distant targets (180 apart) have non-overlapping policies **(D)**, allowing independent decoder adaptations. **(E)** For nearby targets, interference prevents learning: angular errors for the +30 (blue) and −30 (green) rotations fail to decrease, as the decoder cannot simultaneously accommodate opposing rotations. **(F)** For distant targets, the model successfully learns both opposing rotations, with angular errors decreasing toward zero. **(G)** Interference index as a function of angular separation between targets. Interference is maximal when targets are identical (0) and decreases with separation.

We trained our model on this dual adaptation by having an agent that had been trained on two reach targets 180° from each other use the respective policy learned for each target. As with the single target adaptation experiment (Figure 3), only weights in the action decoder network *f* were allowed to change. Consistent with human data, our model learned to correct for visuomotor rotation errors in the version of the task with distinct targets (Figure 4D & F), but could not simultaneously learn two rotations when the initial targets were the same (Figure 4C & E).

We next investigated if there was a relationship between the extent of the generalisation effect and how close targets could be before there was interference in the dual-adaptation experiment. Our model predicts that interference should not be all-or-none, but rather should vary continuously with the angular separation between targets: as targets move closer together in action space, their corresponding policies lie closer in the latent space, leading to greater overlap in the portions of the decoder weights that are modified during adaptation.

To test this, we trained agents on the dual adaptation task with target separations ranging from 0° to 180° in 30° increments. For each separation, we quantified the mean adaptation amount as the absolute angular shift in reaching direction following training. To isolate the effect of interference, we computed an interference index defined as 1 − (*mean adaptation/baseline adaptation*), where baseline adaptation was taken from the 180° separation condition in which the two targets are maximally distinct and thus expected to adapt independently. An interference index of 0 therefore indicates adaptation equivalent to the independent baseline (no interference), while positive values indicate reduced adaptation due to competition between the two rotations. We found that interference decreased monotonically with angular separation (Figure 4G). When targets were identical or separated by only 30°, interference was substantial. As separation increased beyond 90°, interference diminished markedly, and targets separated by 150° or more showed near-complete preservation of adaptation capacity. Together with the generalisation results (Figure 3), these findings suggest a unified account: the same local structure in embedding space that enables generalisation to nearby targets also imposes limits on the capacity to simultaneously learn conflicting adaptations for similar actions.

## Discussion

While it has long been appreciated that reinforcement and supervised learning may serve different functions in the brain [73–75], there are few accounts for the role of interactions between these two learning systems. In this paper, we introduce a novel model of how the brain may use supervised learning of an action representation to aid reinforcement learning in large action spaces. We propose that the cortico-cerebellar system plays the role of supervised network predicting which action causes each state transition, via embedding this mapping in a low-dimensional space. The basal ganglia, in turn, learn a policy over this structured low-dimensional embedding space using reinforcement learning. We show that this model recapitulates the organisation of neural activity in mouse basal ganglia input during reaching movements, the generalisation effects seen in humans performing motor adaptation tasks, as well as their limits when participants have to adapt to multiple perturbations to the same action.

### Relationship to previous models

#### Relationship to previous reinforcement learning models of the basal ganglia

In terms of architecture and learning rule, our model of the basal ganglia is essentially an actor-critic architecture – a popular model of the basal ganglia [12, 60, 76–79]. This means that many of these model’s predictions hold in our formulation. Yet, our model differs from previous reinforcement learning models in that we embedded it within a specialised motor encoder-decoder network that learns low-dimensional action representations. While there has been considerable interest in understanding the neural correlates of reinforcement learning in the brain, many of these focus on learning representations of state, rather than action. For example, theoretical work suggests that the hippocampal formation learns predictive representations of *states* that are useful for reinforcement learning [80–86]. Here we explicitly learn representations of *actions*. State representation learning may be most critical when planning occurs over abstract action spaces – such as discrete choices or movement directions in navigation – where the challenge lies in learning the structure of the environment. In contrast, action representation learning may become essential when learning at the level of motor commands, where the action space is higher-dimensional –think of the number of potential movements, muscle activity patterns, etc. Note that our action representation model and these state representation models are not mutually exclusive: in principle, any type of state representation could be fed into our model. Another possibility is that a reinforcement learning system based on predictive state representations in the hippocampus and an actor-critic-like system in the basal ganglia exist in parallel and compete for control [87].

There is strong experimental evidence that humans and animals are able to learn [1, 88] and adapt [89] movements using reward-based feedback, and reinforcement learning models have been used to explain learning in motor tasks [75, 90–93]. We add to this literature by showing how reinforcement learning can benefit from sensorimotor-error driven learning, modelling these forms of learning not as distinct processes but as a unified synergistic learning system. Furthermore, we show that learning based on sensorimotor errors is also compatible with our model, without needing to alter weights in the reinforcement learning system through slow reward-based updates.

#### Relationship to previous models of supervised learning in the cortico-cerebellar network

Traditionally, motor adaptation has been thought as a process of updating (and then inverting) a “forward model” in which the brain learns to predict the sensory consequences of a given action [30, 31, 33, 94, 95]. The brain then solves the problem of which action to take to reach the desired goal (a future state) – according to this view – by inverting the forward model to calculate an “inverse model” that specifies the right action. Although there are diverse views about the details of their implementation [96–99], the general consensus is that these processes are implemented by the cerebellum. In stark contrast to this traditional view, a recent behavioural study [100], further supported by our modelling work [35], suggests that motor adaptation is best described based on directly updating the control policy (i.e., the inverse model), without the need to invert an updated forward model. In our model, the supervised learning system predicts the action that would cause a particular state transition, bearing strong similarity to a classic inverse model, and thus most strongly aligns to this direct control policy update view. This inverse-like model is not used to control behaviour –we have abstracted away this process – but rather to learn a useful action representation, with behaviour being governed by the interaction between the action decoder and policy networks.

Whether the cortex and cerebellum are both simultaneously performing supervised learning [101, 102], or the cerebellum alone is a supervised learner whose activity shapes that of the cortex [20, 39, 40], remains an open question. Our model is compatible with either view: the cerebellum could learn the action embedding space independently and use it to shape motor cortical dynamics, which would then in turn shape the reinforcement learning process via its direct projections to the striatum.

#### Relationship to other models of corticocerebellar-basal ganglia interactions

Interactions between the cerebellum, cortex, and basal ganglia have been discussed in theoretical terms, but few studies have implemented these ideas as working computational models. Previous ideas have suggested that the basal ganglia serve as a “tutor” for motor cortex [103, 104], or that, conversely, motor cortex tutors the basal ganglia [105, 106]. These tutoring models propose that skills are initially learned in one region and then “transferred” to another as a form of long term storage, much like the reinforcement learning system copies the policy learned by the supervised system in [107]. Other groups have made yet another proposal for how such transfer occurs: that the cerebellum refines coarse motor plans initially learned by the basal ganglia [5, 108]. Unlike these frameworks, our model does not use the multiple brain regions/learning algorithms to transfer learning over time, but rather uses the supervised system (which we propose is analogous to the corticocerebellar network) to learn better representations for reinforcement learning in the basal ganglia. This structure recapitulates key features of basal ganglia activity, and enables rapid updating of the output of a learned policy solely through remapping the action representation to action output transformation, which provides support for its biological plausibility.

### Experimental predictions and future work

We have shown above that our model is consistent with several neural and behavioural findings in the literature. In particular, our model explains why striatal activity patterns are similar for similar movements [54] yet distinct for distinct choices [109], and both observations follow directly from learning policies in a structured action embedding space. It is also consistent with the finding that adaptation with only reward feedback is more prone to noise than adaptation with sensory feedback, and that reward feedback leads to a narrower generalisation curve [89].

Our model also makes a novel prediction: training subjects in a paradigm where dissimilar actions lead to similar sensory consequences should make striatal activity for those actions more similar. This follows from the fact that action embeddings are structured by their consequences: if two motor commands produce similar state transitions, they should occupy nearby regions of embedding space, and thus evoke similar striatal activity patterns. A corollary is that revaluation should generalise more strongly across actions with similar striatal activity than across actions with similar kinematics

Although we have examined three different behavioural tasks, future work should further test the specific predictions of our model regarding what processes are implemented by which regions, and explore how similar principles might apply to other domains that involve higher-dimensional action spaces.

### Conclusions

We have introduced a novel computational model of cerebellar-cortical-striatal interactions during motor learning. In this model, the cortico-cerebellar network uses supervised learning to train an action embedding and a basal ganglia network learns policies in this embedding space. Our model connects several independent neural and behavioural findings, such as the structure of neural activity for similar actions in striatum, the ability to quickly adapt to visuomotor rotations, and particular patterns of generalisation observed in visuomotor rotation experiments. More broadly, our work suggests how the brain’s remarkable ability to learn complex motor skills may arise from the interaction between supervised and reinforcement learning systems.

## Methods

### Formal setting

We model motor learning as a reinforcement learning problem in which agents interact with an environment through a sequence of states (*s* ∈ 𝒮), selecting actions (*a* ∈ 𝒜) to maximise reward. When the agent executes *A*_*t*_ at time *t*, the agent transitions from state *S*_*t*_ to state *S*_*t*+1_ and receives a scalar reward signal, The agent’s aims to find the course of actions, or policy, that maximises the discounted cumulative reward, *G*_*t*_:

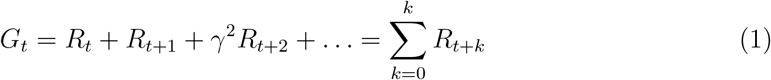

where *γ* ∈ [0, 1] is the discount factor that determines how valuable future rewards are considered. The policy, *π*, is a conditional distribution over actions for each state: *π*(*a*|*s*) := *P* (*A*_*t*_ = *a*|*S*_*t*_ = *s*). The agent’s goal is to find the optimal policy, which maximises the expected discounted cumulative reward, or value, over all states and actions. State value is defined as 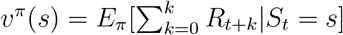 and state-action value as 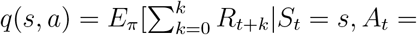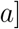.

Reinforcement learning algorithms can be broadly categorised by how they approach the policy optimisation problem. Value-based methods learn the state-action value function *q*(*s, a*) and derive the policy by selecting the highest expected value in each state, *π*(*s*) = *argmax*_*a*_*q*^*π*^(*s, a*). Policy gradient methods take a different approach and directly parameterise the policy, *π*_*θ*_(*a*|*s*) using gradient ascent to directly optimise the policy in terms of expected return. Actor-critic methods combine both approaches: the “critic” learns a value function to estimate expected returns, while the “actor” uses this value estimate to update the policy parameters directly. Actor-critic architectures have frequently been proposed as a model of brain function [76–79] because reward prediction errors, signalled by dopamine [12], can provide the teaching signal to update both the actor (policy) and critic (value function) components. However, all of these methods struggle with high-dimensional action spaces, because of exploration and generalisation challenges. The exploration problem arises because randomly sampling actions in high-dimensional spaces is inefficient. The generalisation problem occurs because these methods lack mechanisms to exploit similarity structure in the action space, treating each action as largely independent.

### Model description

In real-world domains faced by humans and animals, the number of actions | 𝒜 | is often very large. To understand how the brain can learn in high-dimensional action spaces, such as motor control, we implement a model which learns an embedding, *E*, of actions, which is of dimensionality smaller than | 𝒜 |. This embedding allows generalisation of value between actions that cause similar state transitions. The main idea is that the supervised network learns this embedding, and the reinforcement learning network outputs its policy in that space. Here, we will first describe how the embedding is learned by a supervised learning module, and then how the reinforcement learning module learns policies directly in this embedding space.

#### Supervised learning module

The supervised learning module consists of an encoder (*g*) and decoder (*f*) network, which take as inputs the current and successor states, *s*_*t*_ and *s*_*t*+1_, and is trained to predict a stochastic estimate of *P* (*a*|*s*_*t*_, *s*_*t*+1_), the probability that action *a* caused the transition to *s*_*t*+1_ by comparing the predicted action distribution with the action actually taken (Figure 1D). Specifically, this network was trained to minimise the cross-entropy loss:

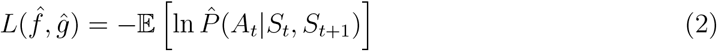

As we will show in the next section, in which we describe the reinforcement learning module, *g* will output an embedding in a space which the policy network will use to learn a policy. The decoder *f*, in turn, projects this action embedding back into the space of environment actions. We chose a one-layer network with tanh nonlinearity for *g* and a single linear layer for *f*. The tanh nonlinearity was chosen so that the action embeddings are bounded, see [43]. Note that, while in principle, *g* and *f* can take the form of any function approximator or neural network, for computational efficiency it is preferable to keep *f* as simple as possible, since it is used both in initial embedding learning and for decision making (see below), whereas *g* is only used for representation learning and can be frozen during decision making as long as the transition probablities *P* (*s*^*′*^|*s, a*) remain constant. The embedding network weights in *g* were initialised from a normal distribution with variance 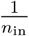, recommended for tanh activations [110]; all other networks used Kaiming uniform initialisation [PyTorch defaults; 111].

### Reinforcement learning module

Although in principle any type of policy gradient method could work (see [43]), our reinforcement learning network implementation consists of an actor-critic network. The critic learns to predict state values, V, as in standard actor-critic methods. The policy network, instead of learning a policy directly in the environment’s action space, learns an internal policy *π*_*i*_ in the learned embedding space, *E*. This policy is a distribution over embeddings given the state, and during decision making, the agent samples an embedding from this distribution:

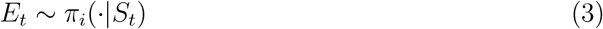

Subsequently, to choose the actual action that the agent takes in the environment, the sample from the policy is mapped through the same action decoder f as the embeddings learned in the supervised network:

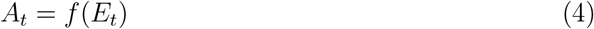

Thus, the policy network learns to output embeddings which are then mapped to actions with the learned action decoder, *f*. The fact that the embeddings in the supervised network and policy network are both mapped through the same action decoder results in policies being learned in the same embedding space as the supervised network. Therefore, in a trained network the policy mean for a given rewarded action will lie close in embedding space to the embedding e that results from inputting the state transition caused by that action into *g*.

In practice, we assume the policy is Gaussian in the embedding space with a standard deviation std. The policy network, a single layer neural network in our implementation, learns to output the mean of this distribution. Embeddings produced by the policy network are also passed through a tanh to keep them in the range of embeddings learned by the supervised network. We anneal the policy standard deviation during learning such that as the reward rate r increases, the standard deviation shrinks (mimicking shrinking uncertainty):

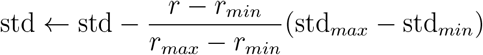

where *r* is the reward rate (average reward over the previous 100 trials), *r*_*min*_ and *r*_*max*_ are the minimum and maximum reward rates used to normalise the annealing schedule, and std_*min*_ and std_*max*_ are the lower and upper bounds on the policy standard deviation (see Table 1).

**Table 1:** Model hyperparameters for all experiments. All experiments were conducted using seeds 0–9. The embedding learning phase (Figure 1) uses LeCun initialisation [110] for *g*. The main learning experiments (Figures 1-2) and adaptation experiments (Figures 3–4) use the pre-trained action embeddings.

| Parameter | Embedding Learning | Main Learning | Adaptation |
| --- | --- | --- | --- |
| *Environment* |  |  |  |
| $d_{action}$ | 24 | 24 | 24 |
| Target angle | — | 135° (2.36 rad) | 135° (2.36 rad) |
| *Reward Structure* |  |  |  |
| radius<sub>max</sub> | 0.5 | 0.5 | 0.5 |
| radius<sub>min</sub> | 0.2 | 0.2 | 0.2 |
| Reward for hit | 1.0 | 1.0 | 1.0 |
| Penalty for miss | −0.1 | −0.1 | −0.1 |
| *Learning Parameters* |  |  |  |
| Number of episodes | 300,000 | 200,000 | 100,000 |
| Actor learning rate $\alpha_a$ | — | $1 \times 10^{-5}$ | 0 |
| Critic learning rate $\alpha_c$ | — | $1 \times 10^{-4}$ | 0 |
| Embedding learning rate $\alpha_e$ | — | 0 | $1 \times 10^{-4}$ |
| Weight decay (embedding) | $1 \times 10^{-4}$ | 0 (frozen) | $1 \times 10^{-4}$ |
| Adam $\beta_1, \beta_2$ | 0.95, 0.999 | 0.9, 0.999 | 0.9, 0.999 |
| Discount factor ($\gamma$) | — | 0.99 | 0.99 |
| *Model Architecture* |  |  |  |
| Embedding dimension ($d_{emb}$) | — | 2 | 2 |
| Fourier order | 3 | 3 | 3 |
| Weight initialisation | $\mathcal{N}(0, 1/n_{in})$ | — | — |
| *Policy Parameters* |  |  |  |
| Inverse temperature ($\beta$) | — | 0.8 | 2.5 |
| Policy std (max) ($std_{max}$) | — | 0.8 | — |
| Policy std (min) ($std_{min}$) | — | 0.2 | 0.2 |
| Reward rate max ($r_{max}$) | — | 0.4 | — |
| Reward rate min ($r_{min}$) | — | −0.1 | — |
| Policy noise ($\sigma_{policy}$) | — | 0.008 | 0.008 |
| *Adaptation-Specific* |  |  |  |
| Visuomotor rotation angle | — | — | −30° |

### Simulation details

#### General centre-out reach task description

We model the centre-out reaching task as a single step reinforcement learning environment where, on each trial, the agent starts at a location in the middle of the environment. There are 24 available actions (length 1) at equally spaced angles. Each action deterministically causes a state transition to one of 24 reach endpoints in the environment (Figure 1C). For simplicity we model the state representation of each reach location using a third order Fourier Basis. This uniquely identifies each possible task state, whilst ensuring that similar reach endpoints have more similar state representations, a property that is critical to learn meaningful embeddings. When the agent makes a reach, it receives both sensory information of the target endpoint and a scalar reward signal, which is 1 if the reach is within a reward zone and -0.1 if not.

The reward zone is a circle whose radius decreased as the reward rate, *r*, increased. The reward rate is the average reward in the previous 100 trials.

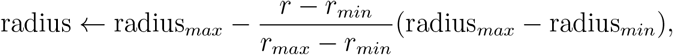

where radius_*min*_ and radius_*max*_ are the lower and upper bounds on the reward zone radius (see Table 1).

#### Training details

The agent’s objective is to find an overall policy, *π*, that maximises the expected cumulative reward. This means training not only the actor and critic networks, but also the supervised module, in order to learn an embedding that the actor network can output its internal policy, *π*_*i*_ into. While it is possible for the policy network to learn at the same time as the supervised network, here we first pre-train the supervised network (embedding pre-training) and then allow the reinforcement learning module to learn policies in that space. Note that these separate learning phases are not necessary for this architecture to work (for formal proof see [43]), but they allow us to identify and clarify the independent contributions of each module. From a biological view, the pre-training phase could be viewed as learning to make reaching movements during development and then learning to reach to a rewarded target later on. In motor learning the mapping from state transition to action is usually constant, except in unnaturalistic situations such as visuomotor rotation experiments which we will model later. Therefore, it is reasonable to pre-learn a mapping from state transition to action prior to learning to assign value to any individual action.

For initial embedding pre-training, the agent chooses random actions to equally sample all actions and corresponding state transitions. Actor and critic network weights are frozen during this time. The supervised network is trained to minimise the cross-entropy loss between predicted and actual actions (Equation 2):

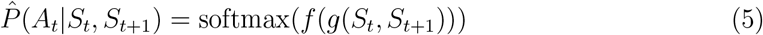

The encoder *g* learns to map state transitions to embeddings, while the decoder *f* maps embeddings back to action probabilities. The loss is backpropagated through both the decoder *f* and encoder *g* networks to learn the embedding representation. We train the supervised network until convergence, after which the encoder and decoder weights are frozen.

In the policy learning phase, we train actor and critic weights, freezing weights in the supervised module. The critic computes the current and next state’s value as 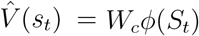 and 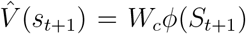, where *W*_*c*_ are the critic’s weights and *ϕ*(*s*) the feature representation of state *s*. It then updates its weights proportionally to the difference between the observed and expected reward:

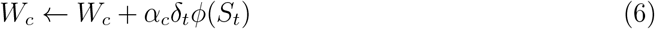

where *α*_*c*_ ∈ [0, 1] is the critic’s learning rate and *δ*_*t*_ = *R*_*t*_ + *V* (*S*_*t*+1_) − *V* (*S*_*t*_) is the reward prediction error. This corresponds to standard temporal difference learning with linear function approximation.

The actor network is updated using a policy gradient method with a baseline. The policy loss gradient is computed as: *L*_*actor*_ = − ln *π*_*i*_(*E*_*t*_|*S*_*t*_) · *δ*_*t*_, where *E*_*t*_ is the embedding sampled from the policy distribution *π*_*i*_(·|*S*_*t*_) and *δ*_*t*_ is the reward prediction error, which serves as an estimate of the advantage.

For embedding pre-training, we trained the supervised network for 300,000 episodes using a learning rate of 1 *×* 10^−4^. The embedding dimension was set to 2.

In the policy learning phase, we trained for 200, 000 episodes with an actor learning rate *α*_*a*_ = 1 × 10^−5^ and critic learning rate *α*_*c*_ = 1 × 10^−4^. Action selection used an inverse temperature parameter of 0.8, which scales the logits before sampling from the policy. We also added Gaussian noise with standard deviation 0.008 to the policy output.

For the adaptation experiments, we trained for 100,000 episodes with an embedding learning rate of 1 × 10^−4^ (with no weight decay) and an inverse temperature of 2.5. All other parameters remained as in the main learning phase. All networks were optimized using the AdamW optimizer [112].

#### Two target reaching task

We extended the single target reaching task to a two target reaching task to simulate the data from Park et al. [54]. As with the single target task there are the same 24 available actions and a third order Fourier Basis provides a state representation of the physical location of the mouse’s paw in space.

While the task that the mice were trained on in Park et al [54] was a reach to pull task, we reasoned that the reaching component can be simplified to a centre out reaching task as we do not simulate the pull phase of the task. As the target is visible to the mice, we reasoned that this visual information can serve as a cue, which we model as the agent learning a different policy mean for each target cue. As with the single target task, we anneal the standard deviation of the policy to the reward rate.

The two targets are 30 degrees apart for the small delta task and 165 degrees apart for the large delta case. While the targets in Park et al. are 10 degrees apart we chose slightly larger spacing for ease of visualisation. The general principle that more similar actions have more similar policies would not be affected by this subtle difference between the experimental and model task parameters.

#### Visuomotor adaptation task

For the single target visuomotor adaptation task (Figure 3), we use the policy network from an agent trained on a single target reach task. In the adaptation phase, actions lead to states that were 30 degrees rotated from the original state they led to. In training the agent to adapt to the visuomotor rotation, all weights in the actor, critic and motor encoder network are frozen and only those in the decoder are allowed to be plastic.

To test the generalisation of the adapted network to other rotated reach targets, we simulate targets being placed at other locations. We simulate there being other targets by placing policy means over the optimal location for a given target, which corresponds to placing the mean of the policy on the point in embedding space that corresponds to the output of g when the state transition for the target action is given as input. To probe for generalisation after the adaptation is learned in the weights of *f* for one target, we freeze weights in *f* after the one target adaptation experiment and sample the embedding space using the simulated policies for the other targets. The angular error reported is the mean of the difference between the angles of the actions taken and the target angle (given the rotation).

For the dual target visuomotor task (Figure 4), we used two learned policies from two different targets but shared a single encoder and decoder network between them. In the extreme case of the targets being on top of each other, the same learned policy was used. Initially one rotation was applied while the agent followed the policy for the corresponding target and the decoder weights changed to account for the perturbation in this direction, then the agent using the already adapted decoder was presented with the other target (and the corresponding policy) with the 30 degree rotation applied in the opposite direction.

All simulation parameters were optimised by hand and are shown in Table 1.

## Acknowledgments

This work was supported by Wellcome Trust (200790/Z/16/Z), the Simons Foundation (564408), ERC MotorAdapt 101169605 and EPSRC(EP/R035806/1). J.A.G. received funding from the EPSRC (EP/T020970/1), Advanced Research + Invention Agency (Scalable neural interfaces SCNI-PR01-P09), and the European Research Council (ERC-2020-StG-949660).

## Author contributions

F.G. and J.P.G. conceived the model, performed simulations, analysed results, and wrote the manuscript. J.A.G. and C.C. contributed to the theoretical framework, model design and links to experimental data, supervised the project, and edited the manuscript. All authors discussed the results and contributed to the final manuscript.

## Code availability

Code to reproduce all simulations and figures will be made available upon publication.

## A Supplementary Figures

**Figure A.1:**
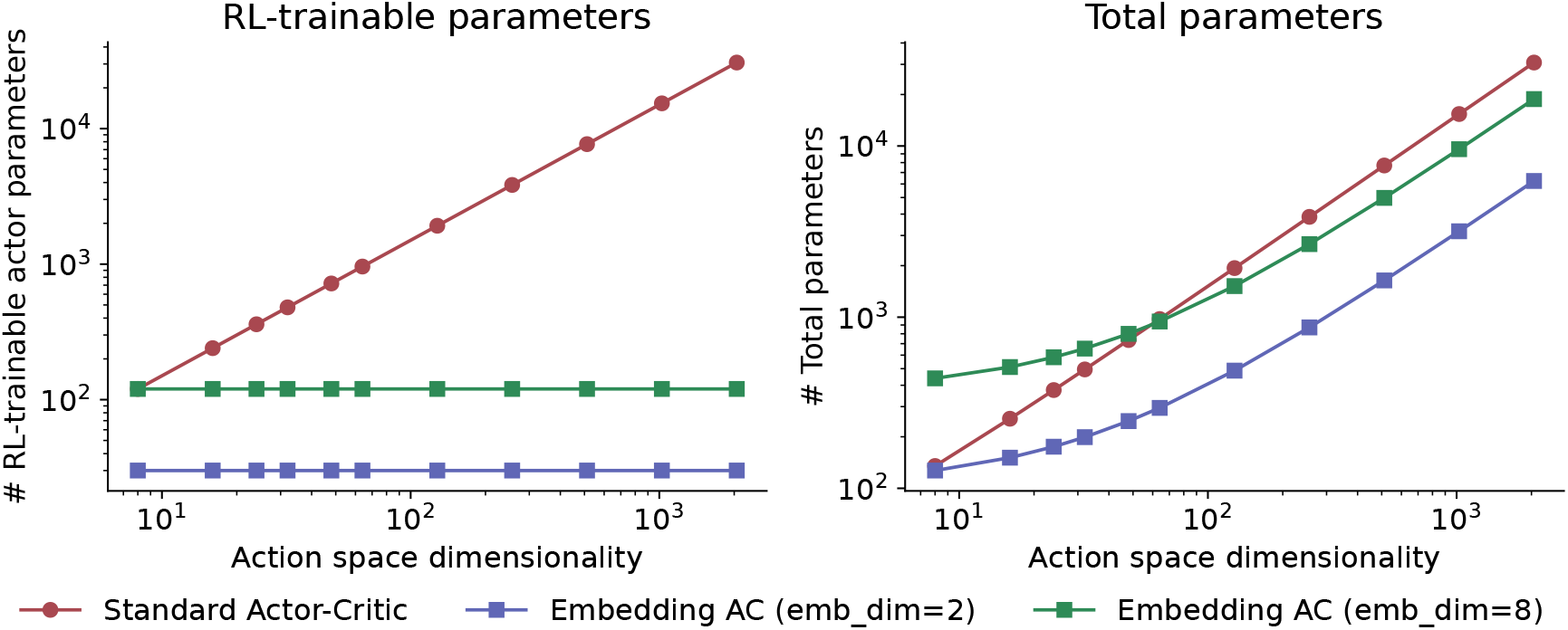
Parameter scaling of standard and embedding-based actor-critic architectures. **(A)** Number of weights (parameters) in the actor network that must be learned during reinforcement learning. For standard actor-critic, the actor maps directly from state features to action logits, scaling linearly with the action space dimensionality (*d*_*state*_ × *d*_*actions*_). For the embedding-based approach, the actor maps from state features to a low-dimensional embedding space, keeping the number of RL-trainable parameters constant regardless of action space size. **(B)** Total parameters including the critic and decoder (*f*) and encoder (*g*) networks used in the embedding approach. While total parameters scale linearly for both architectures, the embedding approach with a 2-dimensional embedding space has a shallower slope (*d*_*emb*_ + 1) × *n*_actions_ vs (*d*_*state*_ + 1) × *n*_actions_). Furthermore, as described in the Methods, the *f* and *g* networks can be pretrained and frozen, amortising this cost across multiple tasks or adaptation scenarios.

**Figure A.2:**
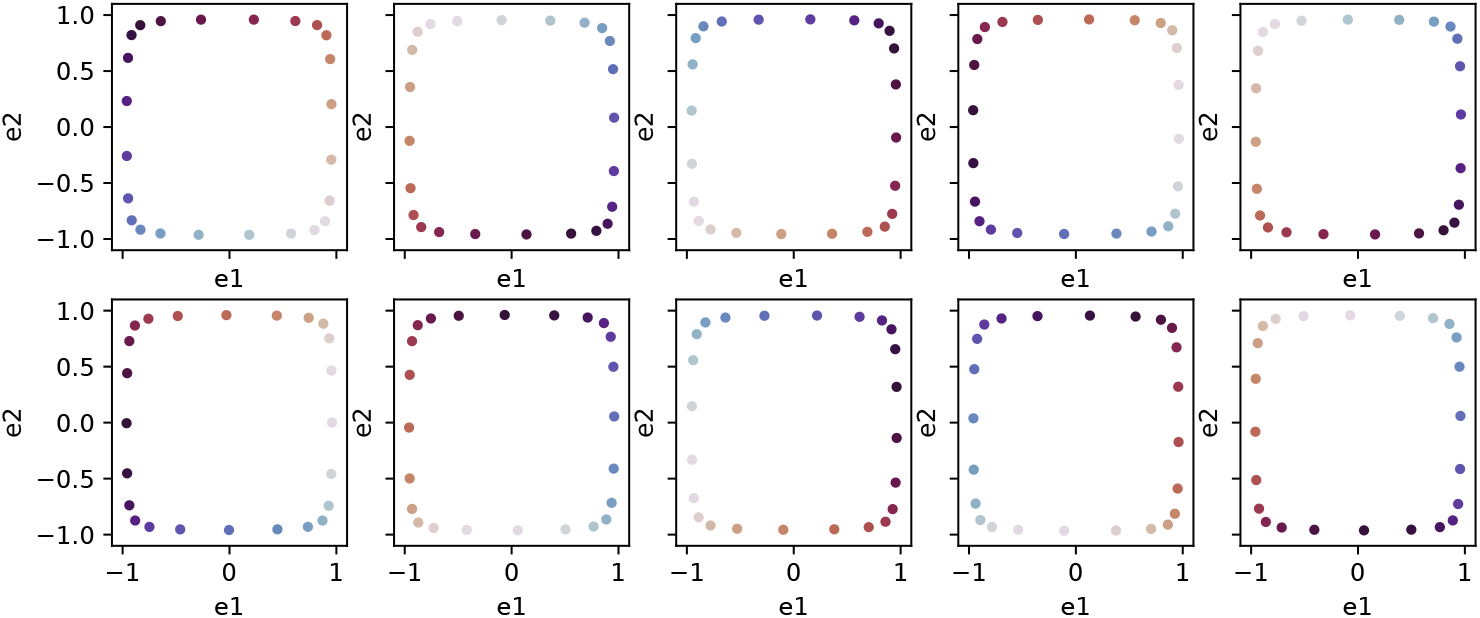
Learned embedding space *e* for random seeds 0-9. Equivalent of Figure 1H.

**Figure A.3:**
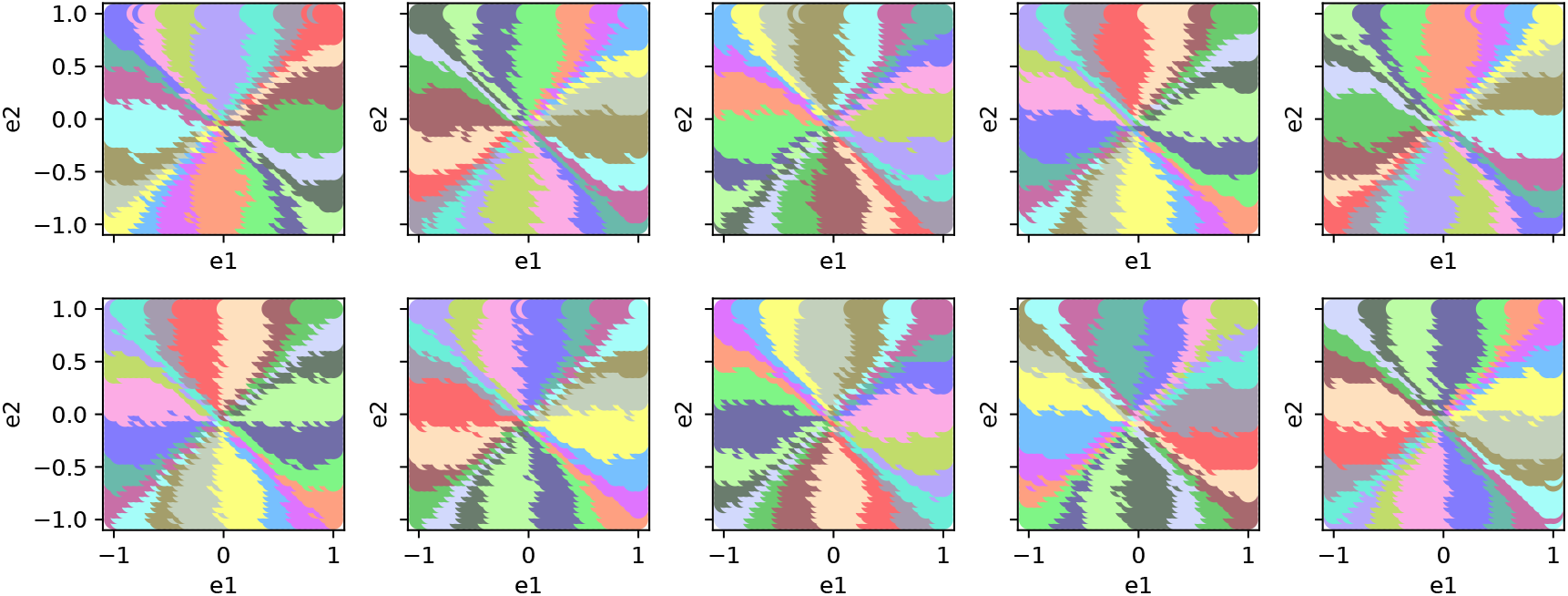
Learned decoder mapping *f* from embedding space to actions for random seeds 0-9. Equivalent of Figure 3G.

**Figure A.4:**
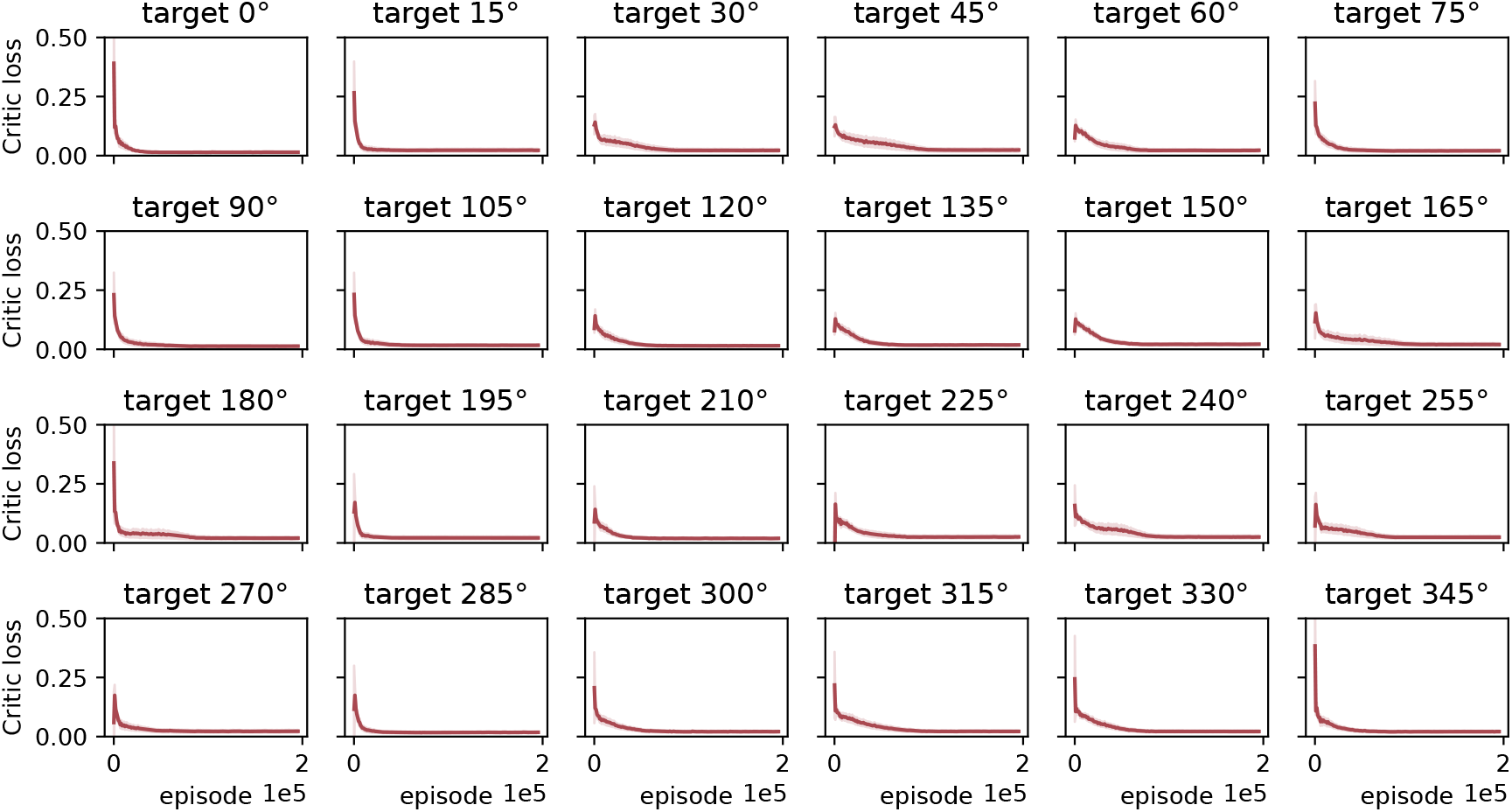
Critic loss (squared TD error) across episodes during reinforcement learning. As in Figure 1I, but for all 24 target directions.

**Figure A.5:**
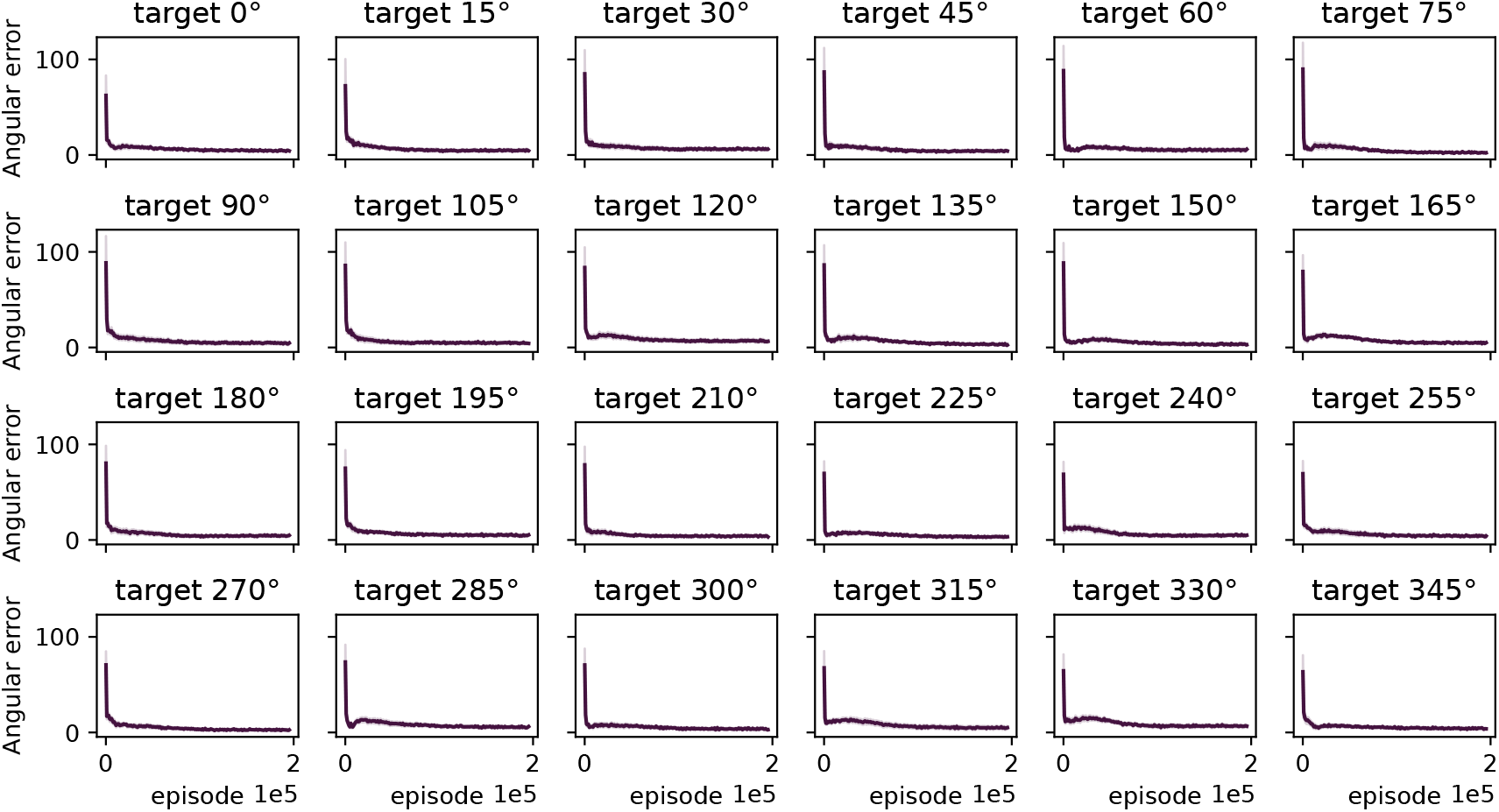
Angular error across episodes during reinforcement learning. As in Figure 1J, but for all 24 target directions.

**Figure A.6:**
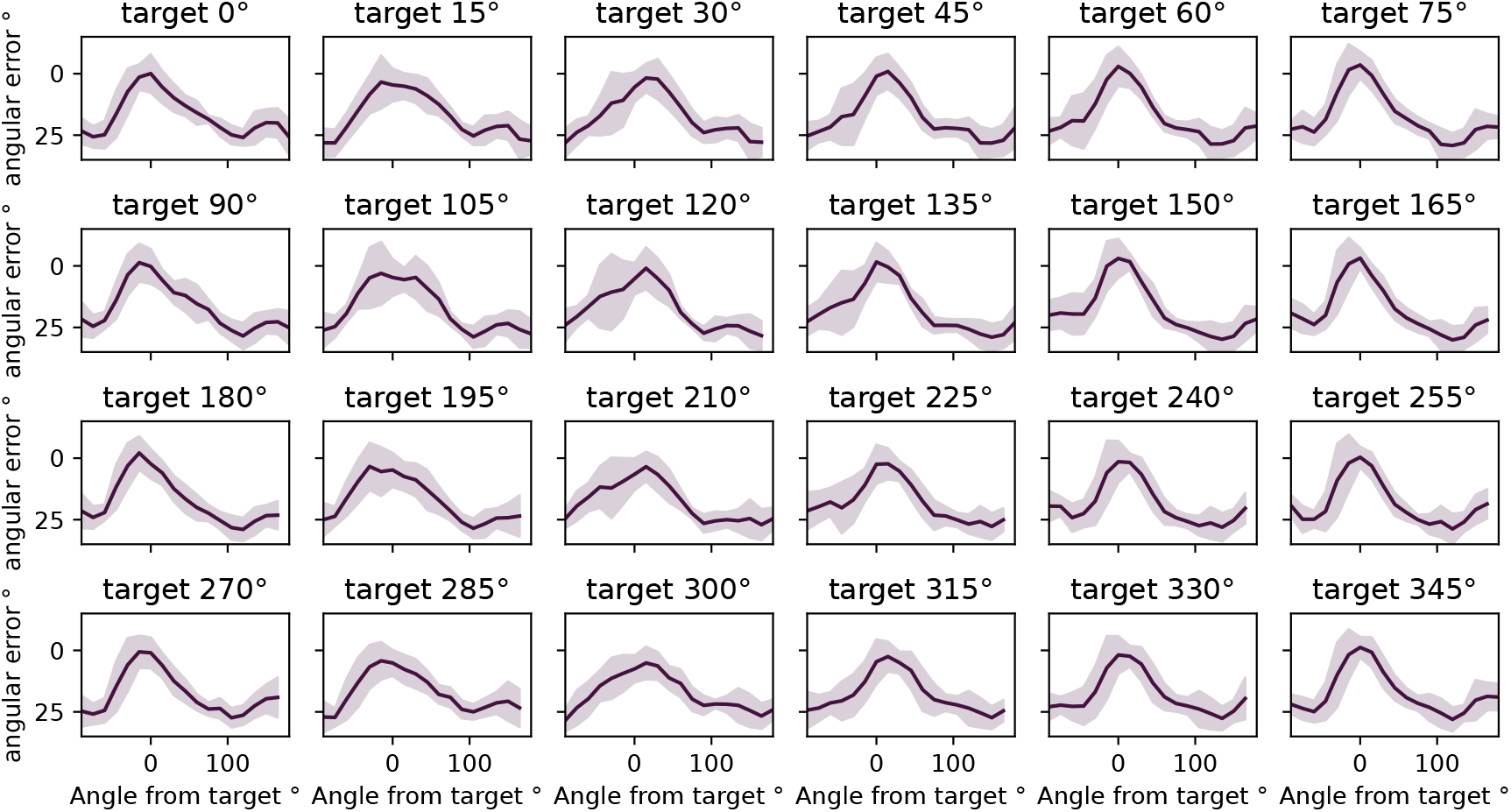
The local adaptation generalisation profile (as in Figure 3I) for each of the 24 initial learning angles.

## Footnotes

1 We use the term ‘representation’ in the machine learning sense [49, 50], unlike the term neural representation [51].

