## Supplementary figures for "Why motor learning involves multiple systems: an algorithmic perspective"

### A Supplementary Figures

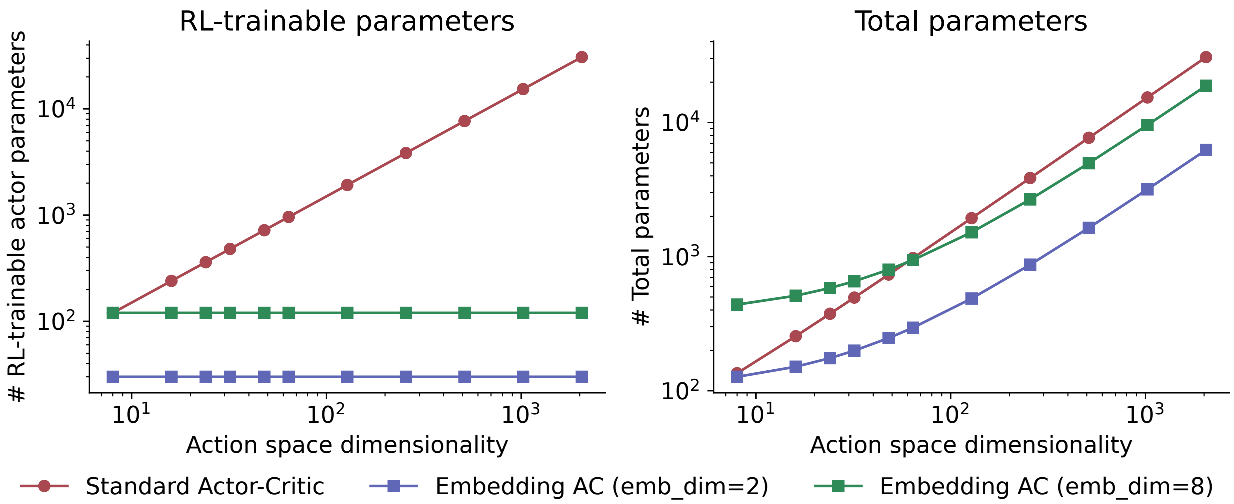


Figure A.1: **Parameter scaling of standard and embedding-based actor-critic architectures.** (A) Number of weights (parameters) in the actor network that must be learned during reinforcement learning. For standard actor-critic, the actor maps directly from state features to action logits, scaling linearly with the action space dimensionality ($d_{state} \times d_{actions}$). For the embedding-based approach, the actor maps from state features to a low-dimensional embedding space, keeping the number of RL-trainable parameters constant regardless of action space size. (B) Total parameters including the critic and decoder ($f$) and encoder ($g$) networks used in the embedding approach. While total parameters scale linearly for both architectures, the embedding approach with a 2-dimensional embedding space has a shallower slope $(d_{emb} + 1) \times n_{\text{actions}}$ vs $(d_{state} + 1) \times n_{\text{actions}})$. Furthermore, as described in the Methods, the $f$ and $g$ networks can be pretrained and frozen, amortising this cost across multiple tasks or adaptation scenarios.

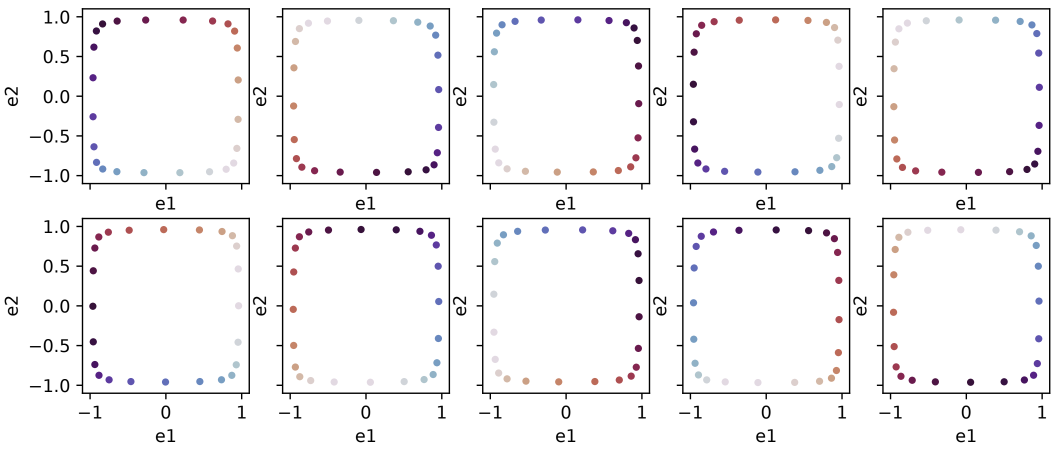


Figure A.2: Learned embedding space $e$ for random seeds 0-9. Equivalent of Figure 1H.

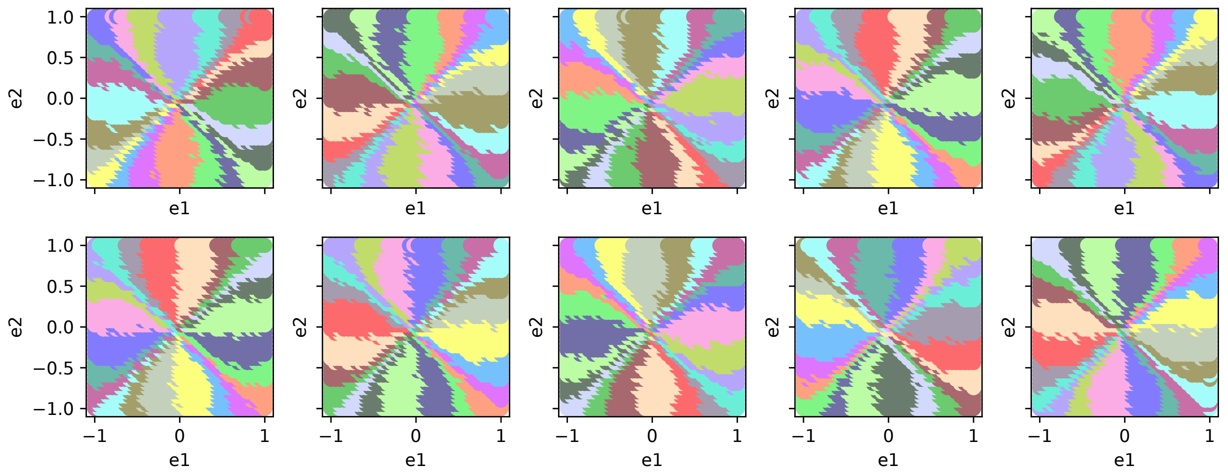


Figure A.3: Learned decoder mapping $f$ from embedding space to actions for random seeds 0-9. Equivalent of Figure 3G.

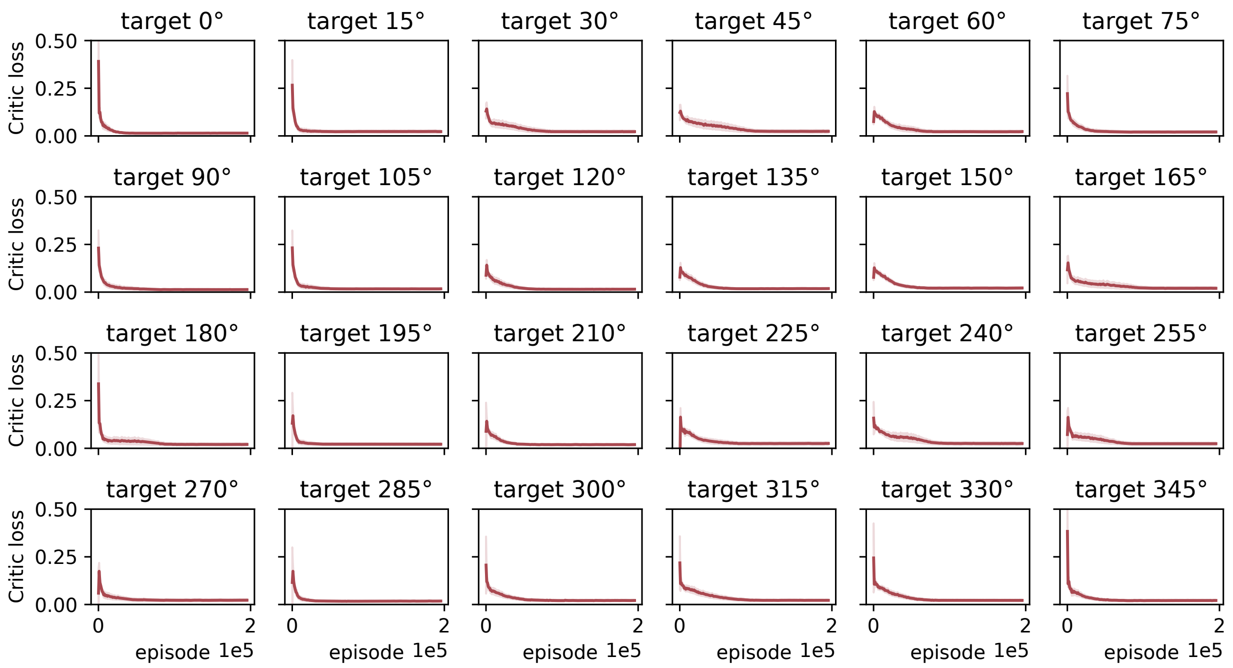


Figure A.4: Critic loss (squared TD error) across episodes during reinforcement learning. As in Figure 1I, but for all 24 target directions.

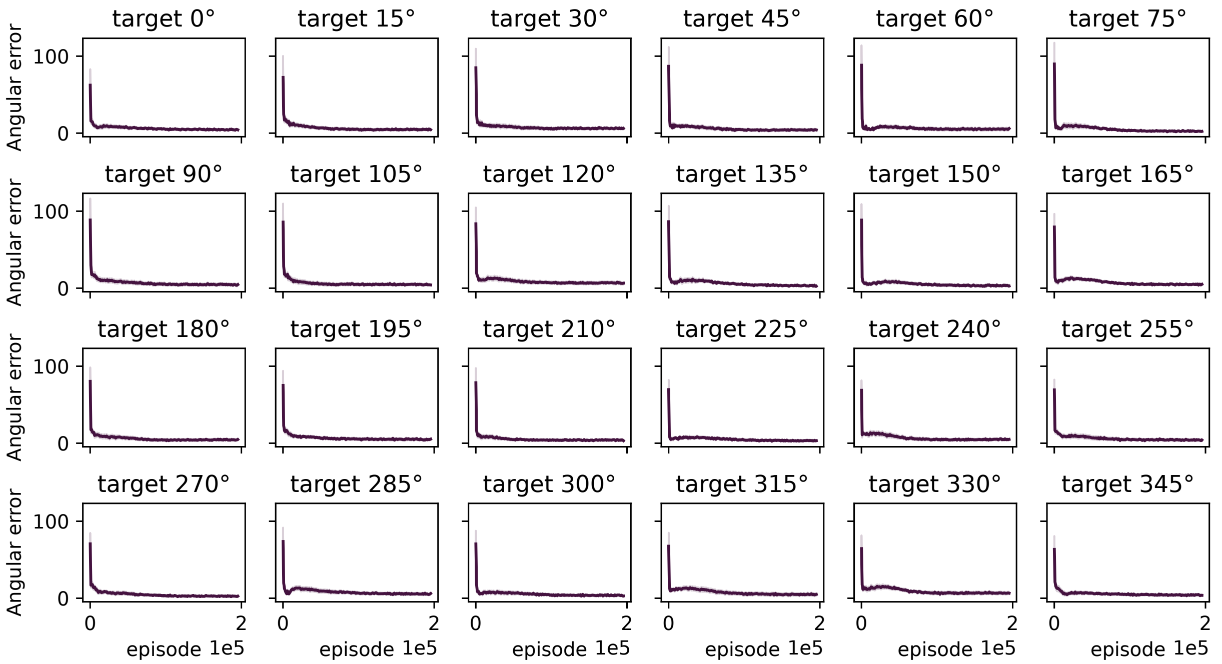


Figure A.5: Angular error across episodes during reinforcement learning. As in Figure 1J, but for all 24 target directions.

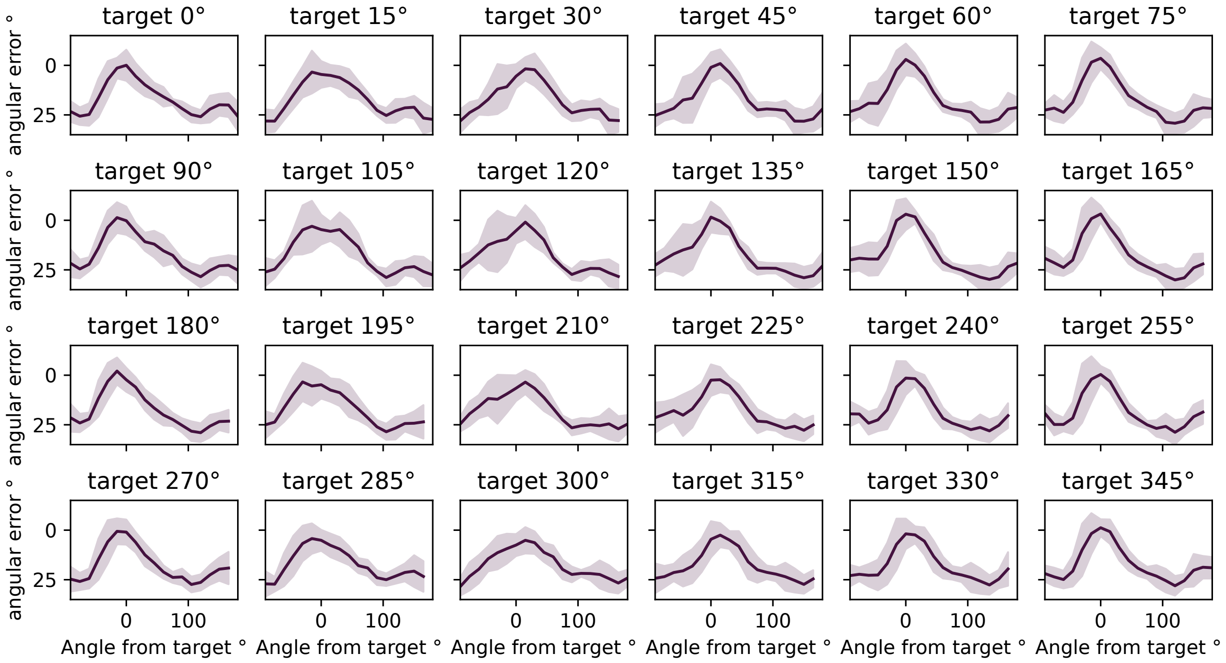


Figure A.6: The local adaptation generalisation profile (as in Figure 3I) for each of the 24 initial learning angles.
